# The water-soluble chitosan derivative, N-methylene phosphonic chitosan, is an effective fungicide against the phytopathogen *Fusarium eumartii*

**DOI:** 10.1101/2021.06.16.448680

**Authors:** Florencia Anabel Mesas, María Cecilia Terrile, María Ximena Silveyra, Adriana Zuñiga, María Susana Rodriguez, Claudia Anahí Casalongué, Julieta Renée Mendieta

**Author notes:** Corresponding authors. E-mail addresses (C. Casalongué), (J. Mendieta).

## Abstract

Chitosan has been considered an environmental-friendly polymer. However, its use in agriculture has not been extended yet due to its relatively low solubility in water. In an attempt to improve such chemical characteristics, a chitosan-derivative prepared by adding a phosphonic group to chitosan N-methylene phosphonic chitosan, NMPC, was obtained from shrimp fishing industry waste from Argentinean Patagonia. This study showed that NMPC had a fungicidal effect on the phytopathogenic fungus *Fusarium solani f. sp. eumartii* (*F. eumartii*). NMPC inhibited *F. eumartti* mycelial growth and spore germination with low IC50 values. *In vivo* studies showed that NMPC affected fungal membrane permeability, ROS production, and cell death. NMPC also exerted antifungal effects against two other phytopathogens, *Botrytis cinerea,* and *Phytophthora infestans*. NMPC did not affect tomato cell viability at the same doses applied to these phytopathogens. Furthermore, the selective cytotoxicity of NMPC could give it added value in its application as an antimicrobial agent in agriculture.

## Introduction

*Fusarium solani* f. sp. *eumartii* (*F. eumartii*) is the causal agent of one of the most economically severe diseases of potato plants. It produces reddish-brown mottling symptoms between leaf veins and dry rot in tubers (Carpenter, 1915). Dry potato rot caused by *F. eumartii* is a threat to several places in the United States, Argentina, Brazil, and Canada. Although *F. eumartii* has been historically considered a potato pathogen, it also infects tomato plants (*Solanum lycopersicum*) (Romberg & Davis, 2007). Some chemical fungicides, such as benzilate and thiabendazole, are commonly used to control fusariosis. However, they are heavy-duty chemicals that often cause detrimental effects, including pollution and toxicity. Specifically, they produce reproductive and developmental problems in laboratory animals at high oral doses. These include skeletal malformations, increased mortality (rats), and multiply anomalies (mice), among others (Gupta, 2018). The resistance of different *Fusarium* spp. has also been described against several chemical fungicides (Hou *et al*, 2018; Qiu *et al*, 2014; Zhou & Wang, 2001). From all these issues, there is an immediate demand for a more sustainable and eco-friendly type of agrochemicals. In this sense, chitosan not only possesses these beneficial characteristics but also does not present toxicity for the environment (Malerba & Cerana, 2018; Maluin & Hussein, 2020).

Chitosan is a linear polysaccharide composed of randomly spread β-(1-4) linked D-glucosamine and N-acetyl-D-glucosamine. This polymer usually comes from chitin, which is abundant and easy to isolate from crustacean exoskeletons. (Younes & Rinaudo, 2015). Most of the agricultural applications reported for the chitosan relates to its capacity for the stimulation of plant defense mechanisms (El Hadrami *et al*, 2010; Hidangmayum *et al*, 2019). Several phytopathological studies demonstrated the antimicrobial properties of chitosan against fungi (Deepmala *et al*, 2015; El Hadrami *et al.*, 2010; Terrile *et al*, 2015), viruses, and bacteria (Badawy *et al*, 2014; Chirkov, 2002; Mania *et al*, 2019; Mansilla *et al*, 2013). However, one disadvantage is that chitosan has a poor-water solubility, so this limitation has restricted its use in agriculture (de Oliveira Pedro *et al*, 2013). The derivatization is the widest procedure used to improve the physicochemical properties, such as solubility (Verlee *et al*, 2017). For that purpose, the production of *O*-, *N*- or *N, O*- substituted derivatives have been extensively employed (Argüelles-Monal *et al*, 2018). Previously, Heras et al. (2001) have described an N-derivatization process by reacting chitosan with a phosphonic group and named it N-methylene phosphonic chitosan (NMPC). In addition to the fact that NMPC is soluble in water over a wide range of pH values, it is a Ca^2+^ and transition metal chelator (Ramos *et al*, 2003). NMPC also showed improved performance compared to chitosan as a non-viral gene carrier in HeLa cells, indicating its high potential in clinical applications (Zhu *et al*, 2007).

This work aimed to study the antimicrobial effect as well as downstream events associated with the mode of action of NMPC-derived chitosan on the phytopathogen *F. eumartii*. Furthermore, we evaluated the antimicrobial activity of NMPC on two other relevant phytopathogens, *Botrytis cinerea* and *Phytophtora infestans*. In conclusion, these findings provided fundamental knowledge on NMPC as a potential antimicrobial agent for modern agriculture.

## Materials and Methods

### Biological materials

Estación Experimental Agropecuaria (EEA) INTA, Balcarce (Argentina) provided *F. eumartii* isolated 3122, which was maintained on solid potato dextrose agar (PDA; Merck, Germany) medium at 25°C in darkness. Spores were collected from 8-day-old culture plates and suspended in sterile distilled water (Terrile *et al.*, 2015). *Botrytis cinerea* strain B05.10 was cultured as described by Benito et al. (1998). *Phytophthora infestans* mating type A2 was grown and preserved on fresh potato tuber slices, as Andreu et al. (2010) described.

Tomato cell suspensions (*S. lycopersicum* cv. Money Maker, line Msk8) were provided and grown in Murashige-Skoog medium as described by Laxalt et al. (2007).

### NMPC preparation

The preparation of chitin and chitosan, as well as the synthesis of NMPC, was carried out as described by Heras et al. (2001). Briefly, we dissolved chitosan (2% w/v) in 1% (v/v) glacial acetic acid. Equals parts of chitosan and phosphorous acid (w/w) were mixed drop-wise with continuous stirring for 1 hr. Then, we increased the temperature to 70°C, and an equal part of 36.5% (w/v) formaldehyde was added drop-wise for an additional 1 hr with reflux. After that, we kept the incubation at 70°C for 5 hr. The clear pale yellow solution was dialyzed against distilled water in dialysis tubing with a cut-off value of 2500 Da for 48 hr or until the pH of the water was raised to 6.8. Finally, the solution was frozen and freeze-dried. We characterized the NMPC as described by Heras et al. (2001). The characteristics of NMPC used in this study are 615,595 Da, viscosity 22.5 mPa/seg, substitution degree 1.54, elemental analysis (%) C, 34.68; H, 7.10; N, 5.15; P, 7.93. The solubility of NMPC in aqueous media over an extended pH range and its filmogenic nature was verified as previously described. We also performed IR spectroscopy of NMPC, as Heras et al. (2001) described.

### Measurements of spore and sporangium germination

We evaluated the antifungal activity of NMPC on *F. eumartii* and *B. cinerea* spores and *P. infestans* sporangia as described by Mendieta *et al.* (2006)*. F. eumartii* (1 × 10^6^ spores/mL) and *B. cinerea* spores (1 × 10^5^ spores/mL), and *P. infestans* sporangia (5 × 10^4^ sporangia/mL) were treated with different concentrations of NMPC (0.5, 1, 1.5, 2.5, 5, 10 μg/mL) in a final volume of 50 μL of 1% sucrose and put on micro slides.The spores of *F. eumartii* and *B. cinerea* were incubated at 25°C, while *P. infestans* sporangia at 18°C for 24 hr in darkness. Germinated spores and sporangia were quantified under light microscope Eclipse E200 (Nikon, Japan) using a hemocytometer. We considered spores and sporangia germinated when the germ tube length was longer than one-half of the reproductive structure (Plascencia-Jatomea *et al*, 2003). We analyzed at least 250 spores or sporangia per replicate, with 3 replicates per treatment. We estimated the IC_50_ values as the NMPC concentrations that reduce germination by 50%.

### *F. eumartii* mycelial growth inhibition

We added different volumes of NMPC (final concentrations were 5, 50, 100, or 500 μg/mL) and a 0.5 cm-diameter disk of PDA agar containing *F. eumartii* mycelia in flasks with 100 mL of PDB media. *F. eumartti* was grown at 25°C with shaking at 100 rpm in darkness. After four days, we filtered each fungal culture through muslin to get the mycelia and placed them in an oven at 65°C for 3 hr. We measured the mycelial-dry biomass, and we estimated the IC_50_ value.

### Fungicidal activity on *F. eumartii* cells

We incubated *F. eumartii* spores (1 × 10^4^ spores/mL) with 1 and 5 μg/mL of NMPC or distilled water in a final volume of 60 μL. Samples were incubated at 25°C for 24 hr in darkness and then spread on PDA. After three days, we counted the colonies and calculated the number of colony-forming units (CFUs) in each sample.

### Fungal cell viability assay

*F. eumartii* cell viability was determined by propidium iodide (PI; Sigma-Aldrich, USA) exclusion as described by Terrile et al. (2015). PI is used to evaluate cell viability as a nucleic acids stain. Once the dye is bound to nucleic acids, its fluorescence is enhanced 20–30-fold (Novo *et al*, 2000). We treated *F. eumartii* spores (1 × 10^6^ spores/mL) with 1 and 5 μg/mL of NMPC at 25°C for 24 hr in darkness. We added PI at a final concentration of 120 μM, and we observed the *F. eumartii* spores in an Eclipse E200 microscope (Nikon, Japan) with a G-2E/C filter set containing an excitation filter at 540/25 nm, suppressor filter at 630/60 nm, and a dichroic mirror at 565 nm.

### Membrane permeabilization assay

We detected fungal cells with compromised cell membranes by recording the fluorescence of the DNA-binding dye SYTOX green (Molecular Probes, USA). Permeabilization of the fungal membrane allows the dye to cross the membranes and to intercalate into the DNA. This association displays an intense fluorescence emission when it is excited by blue light illumination (Rioux *et al*, 2000). We incubated the spores with 5 μg/mL of NMPC or distilled water at 25°C for 1, 2, and 4 hr in darkness. Next, we added 1 μM of SYTOX Green, and we immediately observed the spores with an Eclipse E200 fluorescence microscope equipped with a B-2 A fluorescein filter set (Nikon, Japan).

### Measurements of endogenous H_2_O_2_

The endogenous H_2_O_2_ level was assessed by a peroxidase dependent staining using 3, 3’-diaminobenzidine (DAB; Merck, Germany). DAB polymerizes in contact with H_2_O_2_ in the presence of peroxidase, producing an insoluble colored complex (Thordal-Christensen *et al*, 1997). *F. eumartii* spores at a final concentration of 1.5 × 10^6^ spores/mL were incubated with 2.5 and 5 μg/mL of NMPC at 25°C for 4hr. Then, 0.5 mg/mL of DAB was added to each sample and incubated for an additional 1 hr before rinsing. We observed the spores under an Eclipse E200 light microscope (Nikon, Tokyo).

### Tomato cell viability assay

Tomato cell suspensions were grown in Murashige-Skoog medium (Duchefa, The Netherlands) supplemented with 5.4 M naphthalene acetic acid, 1 M 6-benzyladenine, and vitamins (Duchefa, The Netherlands) at 24°C with continuous agitation in darkness as described by Laxalt et al. (2007). We tested tomato cell viability by Evans blue staining assay (Sukenik *et al*, 2018). We incubated the cells with 10 and 100 μg/mL of NMPC at 30°C for 24 hr in darkness. As a positive control, we treated cells with 1% Triton X-100. Twenty-five μL of 1% Evans blue solution were added to 50 μL of treated-suspension cells, incubated at room temperature for 5 min, and observed under Eclipse E200 light microscope (Nikon, Tokyo). The extent of dye uptake by dead cells was quantified spectrophotometrically by incubating 250 μL of each suspension with 150 μL 1% Evans blue for 5 min at room temperature. Unbound Evans blue stain was removed by washing four times with 0.1 M Tris-HCl pH 7.5. Cells were collected by centrifuging at 800 rpm for 15 sec and lysed with 250 μL 100% dimethyl sulfoxide (Sigma-Aldrich, USA) at 100°C for 15 min. We measured the absorbance at 595 nm by using a microplate reader ELx800 (BioTek, USA).

### Statistical analysis

The values shown in each figure are the mean values ± SD of at least 3 experiments. Data were subjected to analysis of variance (one-way ANOVA) and post hoc comparisons with Tukey’s multiple range test at P <0.05 level. We used GraphPad Prism 5 (GraphPad Software Inc., San Diego, CA, USA) as a statistical software program.

## Results

### NMPC is an antifungal chitosan derivative

As a first approach to evaluate the antifungal effect of NMPC, we incubated *F. eumartii* and *B. cinerea* spores and *P. infestans* sporangia with different concentrations of NMPC for 24 hr. Inhibition of germination of both cell types by NMPC was dose-dependent, being almost 100% of spores and sporangia inhibited at 10 μg/mL NMPC. (Fig. 1a and 1b). The estimated IC_50_ values for *F. eumartii*, *B. cinerea*, and *P. infestans* were 2.5 ± 0.9, 4 ± 1.2, and 2 ± 1.3 μg/mL, respectively. Besides, the IC_50_ for *F. eumartii* was in the same range as that obtained using chitosan (4.3 μg/mL ± 2.3), which is the precursor of NMPC (Supplementary figure). Next, we studied in depth NMPC action on *F. eumartti* phytopathogen. A significant dose-dependent mycelial growth inhibition was also observed, with a dry mass reduction of nearly 60% at 50 μg/mL of NMPC (Fig. 2). In this case, the estimated IC_50_ for mycelial growth was 22 ± 5.2 μg/mL, nearly nine times higher than the estimated IC_50_ value for *F. eumartii* spore germination.

**Fig. 1.**
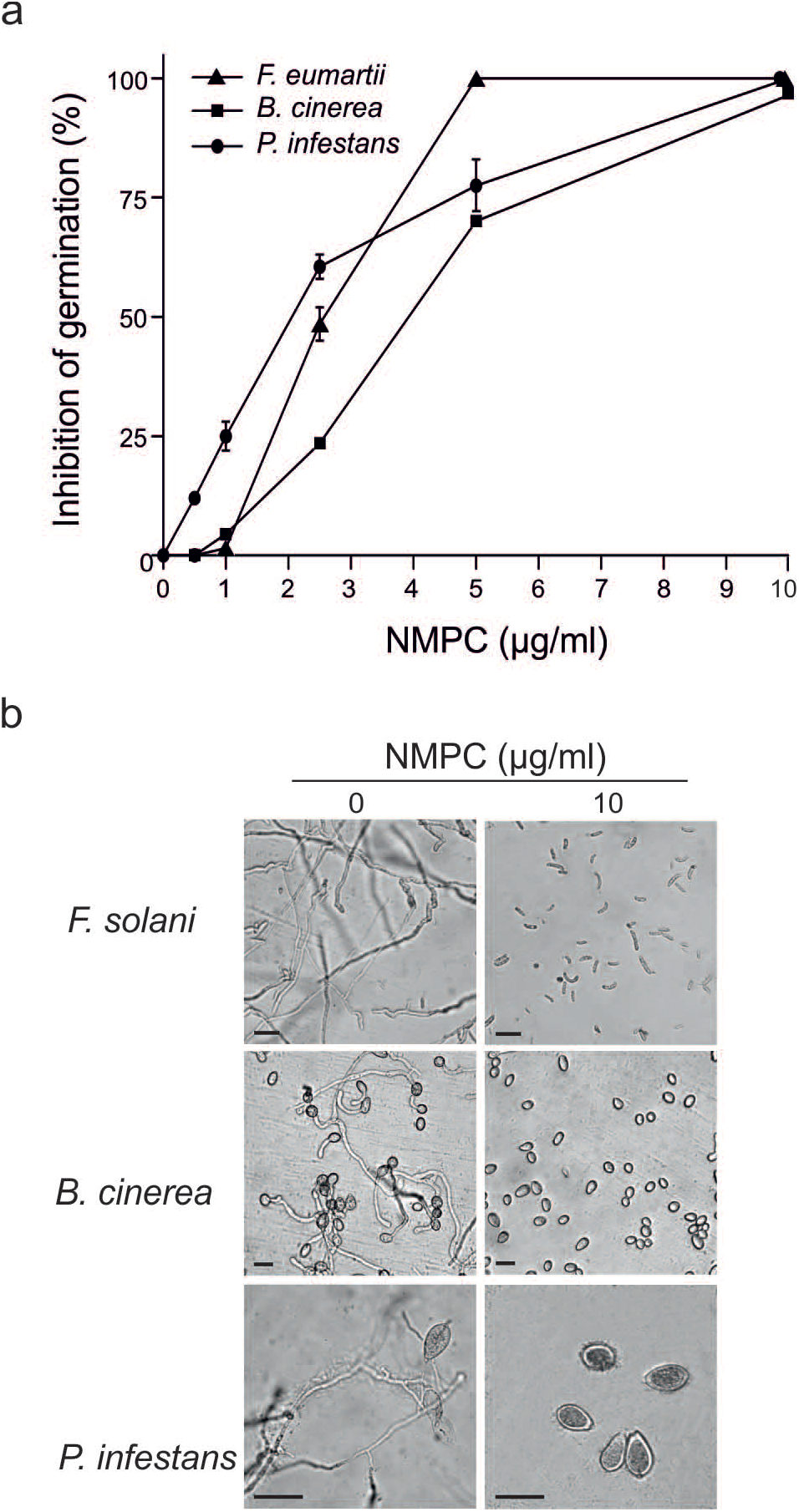
Antimicrobial dose-dependent effect of NMPC. (a) The values represent the percentages of total spores/sporangia present in each sample after the incubation with different concentrations of NMPC for 24 hr. Each value is the mean ± SD of at least 3 independent experiments. (b) Representative images of the spores/sporangia are shown. Scale bar: 22 μm (upper panels), 15 μm (middle panels) and 30 μm (lower panels).

**Fig. 2.**
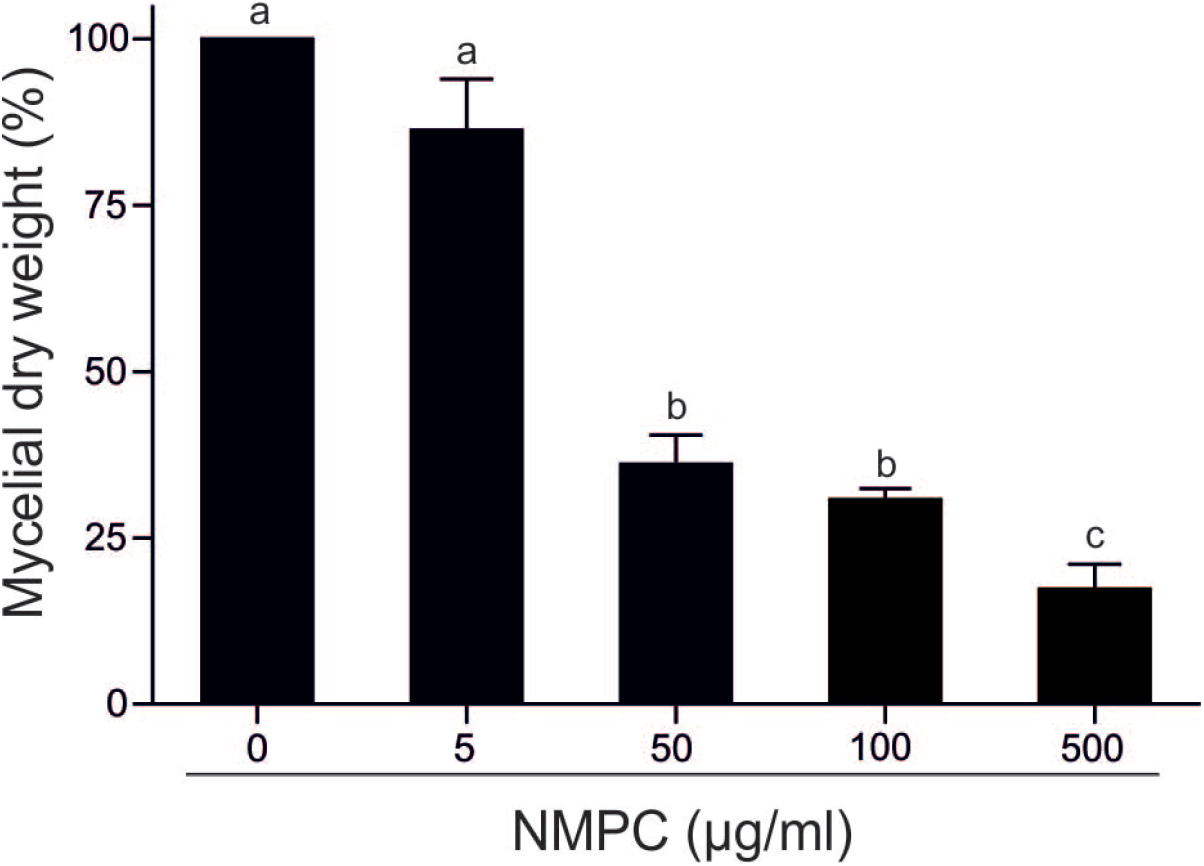
NMPC inhibits *F. eumartii* mycelial growth. *F. eumartii* was inoculated in a liquid PDB medium supplemented with different concentrations of NMPC and incubated for four days. The quantification of the mycelial dry weight is expressed as the percentage of control (100%). Each value is the mean ± SD of 3 independent experiments. Different letters point out statistically significant differences (Tukey’s test, *p* < 0.05).

To study the fungicidal effect, we incubated *F. eumartii* spores with different concentrations of NMPC for 24 hr and then plated on an NMPC-free PDA medium for three days. Interestingly, incubation with 5 μg/mL of NMPC almost completely abolished fungal growth, suggesting an NMPC-mediated fungicidal action (Fig. 3). We also analyzed *F. eumartii* cell viability by PI staining. Only the cells that have damaged plasma membranes take up PI, and the red fluorescence is a consequence of DNA–dye binding. As shown in Fig. 4, while control spores remained unstained, an increase in the percentage of PI-positive spores was observed in NMPC treatment, being higher at a dose of 5 μg/mL. The PI-positive spore percentage was 55.5% and 91.3% for 1 and 5 μg/mL of NMPC, respectively (Fig. 4b).

**Fig. 3.**
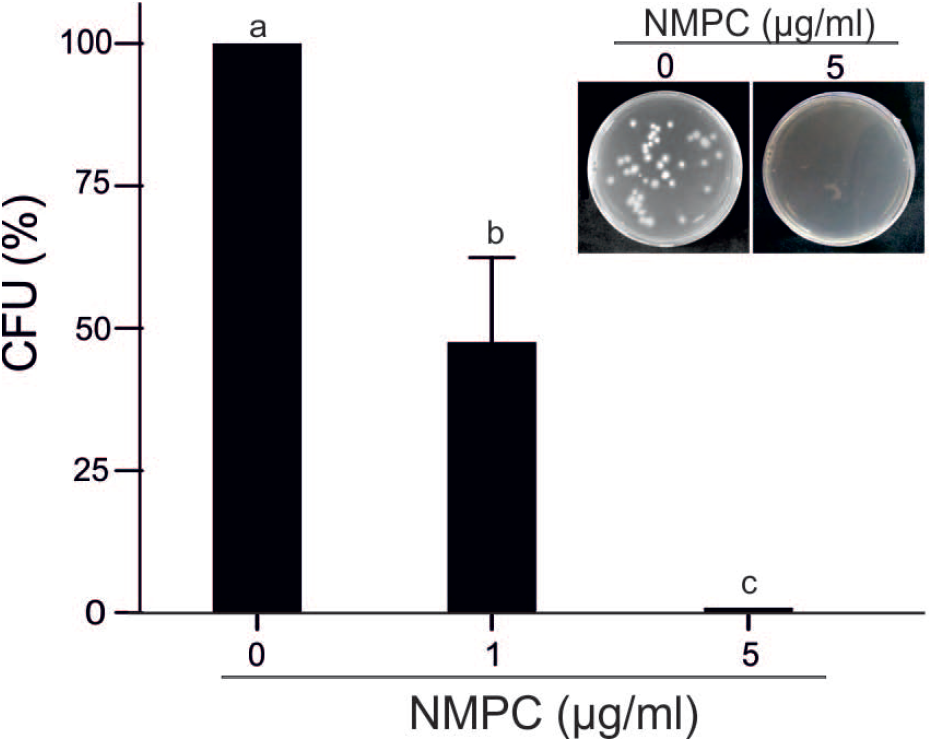
NMPC mediates fungicidal action on *F. eumartii* spores. Spores were incubated with 1 or 5 μg/mL NMPC for 24 hr and then plated on the Petri dishes containing fresh PDA media to allow fungal growth. *F. eumartti* was grown for three days at 25°C. Values represent the percentage of CFU of control (100%). Each value is the mean ± SD of at least 3 independent experiments. Different letters point out statistically significant differences (Tukey’s test, *p* < 0.05). Representative images are shown (inset).

**Fig. 4.**
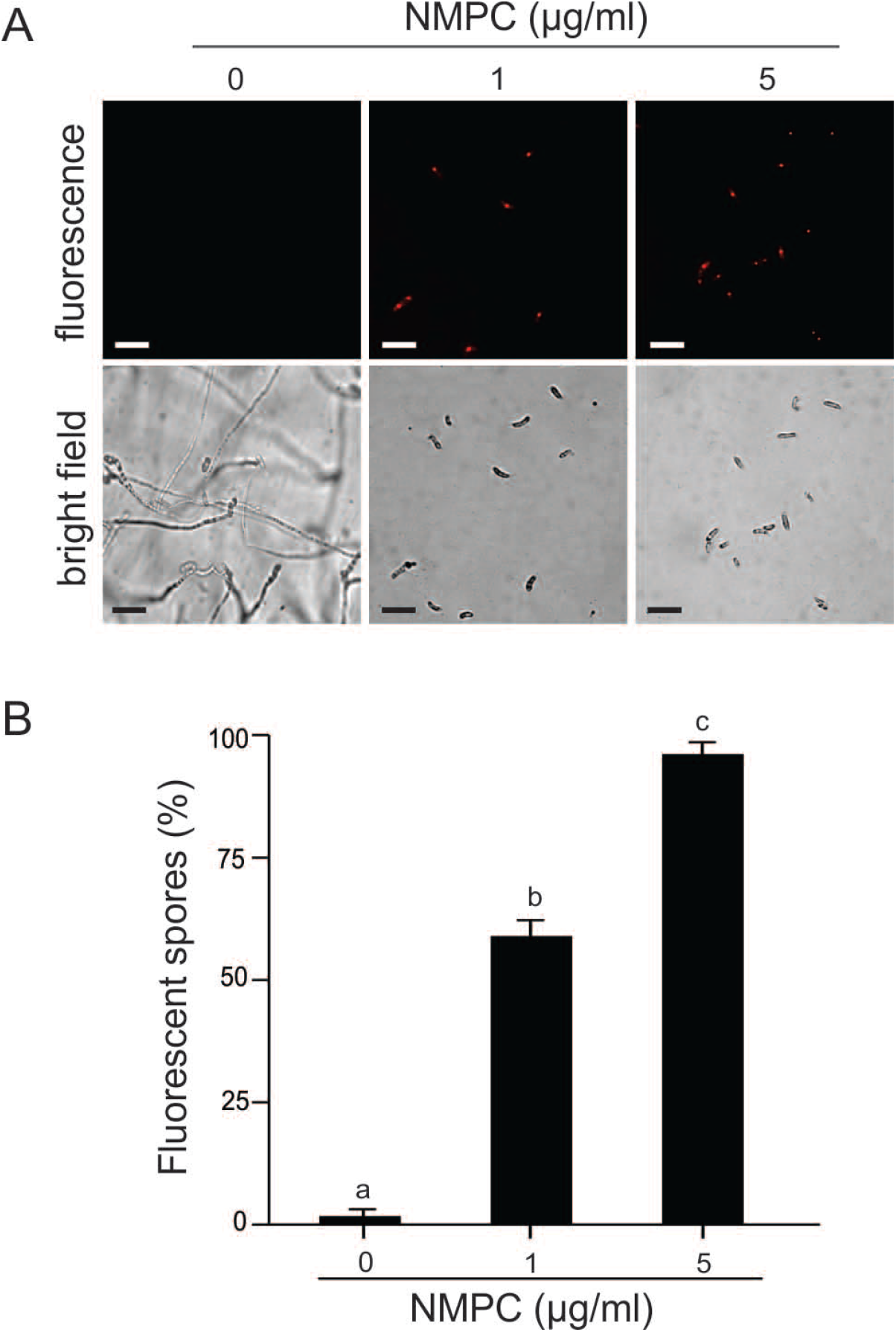
NMPC affects *F. eumartii* cell viability. Fungal spores were incubated with 1 μg/mL or 5 μg/mL NMPC for 24 hr and stained with PI. The dead spores are observed in red fluorescence. (a) Representative images are shown. The bright-field image for each treatment is shown below the respective fluorescent images. (b) Values represent the percentage of the red spores present in each sample. Each value is the mean ± SD of at least 3 independent experiments. Different letters point out statistically significant differences (Tukey’s test, p < 0.05). Scale bar: 22 μm.

### NMPC triggers membrane permeabilization and endogenous H_2_O_2_ in *F. eumartii* cells

The cell membrane integrity of *F. eumartii* spores was analyzed by using SYTOX Green. Spores incubated with 5 μg/mL of NMPC during different times were stained with SYTOX Green and subjected to microscopic analysis. Fig. 5a shows that SYTOX Green-mediated fluorescent spores increased over time. While at 1 hr after NMPC treatment, 22% of spores displayed green-fluorescence, 4 hr after, 65% of them were positive for SYTOX Green (Fig. 5b). Next, NMPC-mediated cytotoxicity was explored by measuring endogenous H_2_O_2_ production in *F. eumartii* spores. The levels of H_2_O_2_ gradually increased in an NMPC dose-dependent manner. After 4 hr of 5 μg/mL NMPC treatment, 98% of spores were stained (Fig. 6). Together, these findings indicated that both cell membrane permeabilization and H_2_O_2_ production could lead to NMPC-induced cell death in *F. eumartii* spores.

**Fig. 5.**
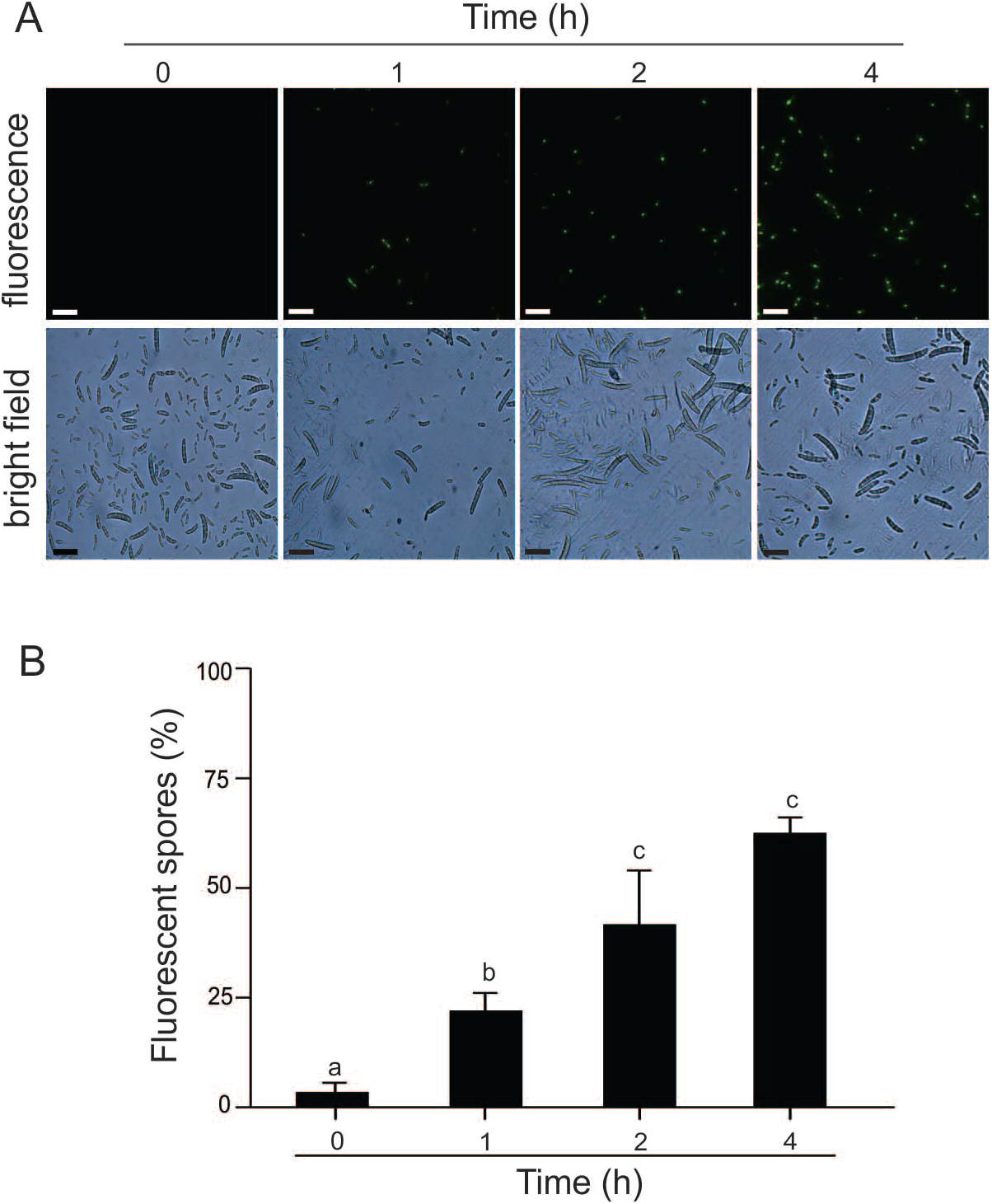
NMPC induces cell membrane permeabilization. Kinetic of cell membrane permeabilization by 5 μg/mL NMPC in fungal spores. Cell membrane permeabilization was visualized in *F. eumartii* spores by the SYTOX Green probe. (a) Representative images are shown. (b) Values represent the percentage of the green-spores present in each sample. Each value is the mean ± SD of at least 3 independent experiments. Different letters point out statistically significant differences (Tukey’s test, p < 0.05). Scale bar: 22 μm.

**Fig. 6.**
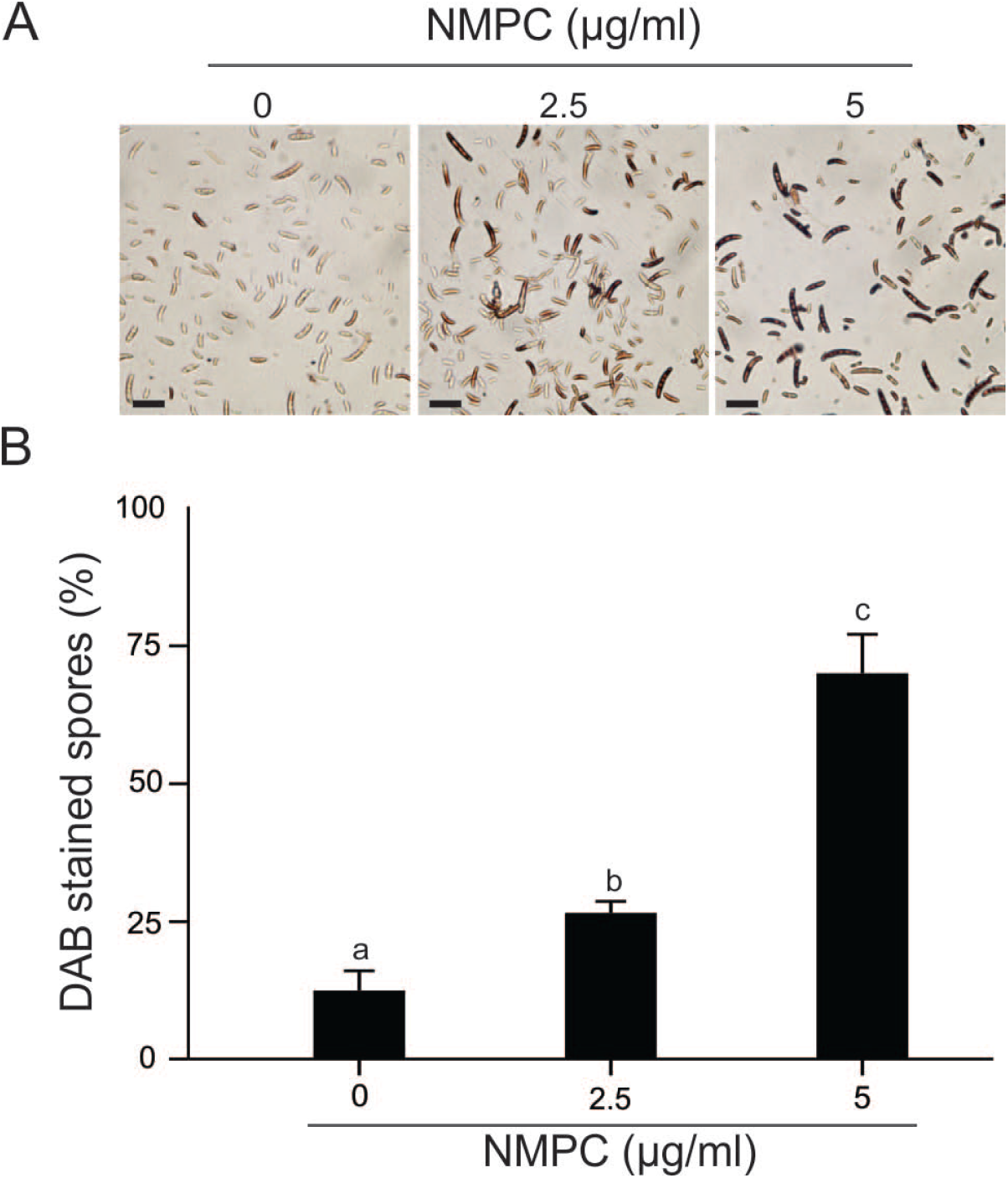
NMPC induces H_2_O_2_ production in *F. eumartii* spores. Spores were treated with 2.5 or 5 μg/mL NMPC for 4 hr before DAB staining and subjected to microscopic analysis. (a) Representative images are shown. (b) Values are expressed as a percentage of total spores in each sample. Each value is the mean ± SD of at least 3 independent experiments. Different letters point out statistically significant differences (Tukey’s test, *p* < 0.05). Scale bar: 22 μm.

### NMPC does not affect tomato cell viability

Considering that we studied NMPC as a putative antimicrobial agent with a projected application on horticulture, we tested its toxicity on tomato cells. Tomato cell suspension cultures were incubated with 10 and 100 μg/mL NMPC for 24 hr and then stained with the vital dye Evans blue. Quantification of dye uptake showed that cell viability did not significantly decrease with 10 and 100 μg/mL NMPC (Fig. 7b). Most of the tomato cells were unstained after 10 μg/mL NMPC treatment (93% ± 6). Even at 100 μg/mL NMPC addition, tomato cells excluded the dye (85% ± 19), and their morphology remained unchanged (Fig. 7a). However, control cells treated with 1% Triton X-100 showed nuclei and cytoplasm fully stained, and the cell size look liked smaller than those treated with water as well as NMPC (Fig. 7a). Cell viability after Triton X-100 treatment was 7 % ± 6 (Fig. 7b).

**Fig. 7.**
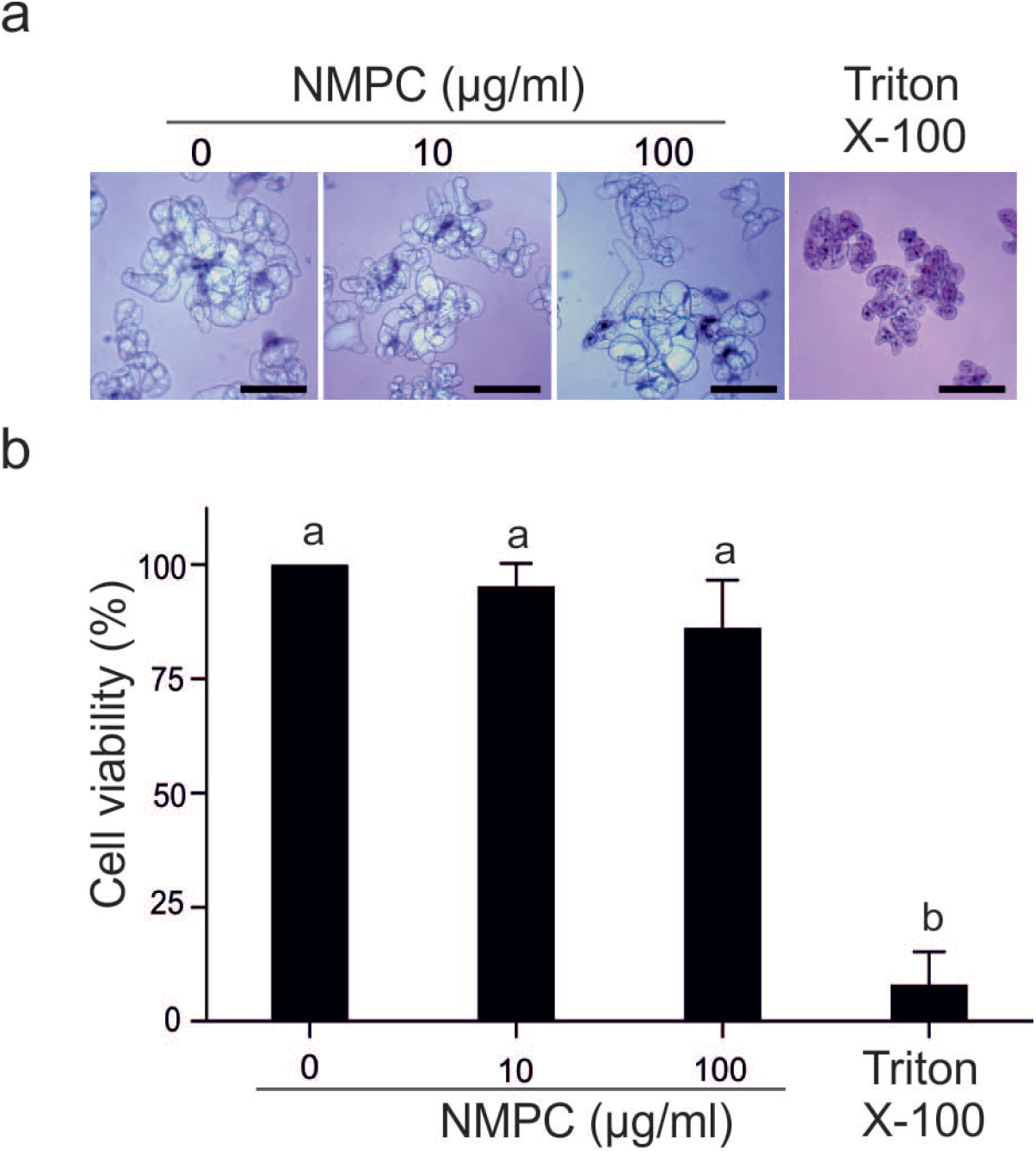
NMPC does not affect tomato cell viability. Suspension-cultured tomato cells were incubated with different concentrations of NMPC for 24 hr and then stained with Evans blue. As negative and positive controls, water and 1% Triton-100 were used, respectively. (a) Representative images of at least 3 independent experiments are shown. (b) Quantification of cell viability was estimated by recording Evans blue retention in tomato cells. Values are expressed as the percentage of water treatment (100%). Each value is the mean ± SD of at least 3 independent experiments. Different letters point out statistically significant differences (Tukey’s test, *p* < 0.05). Scale bar: 20 μm.

## Discussion

In this study, we demonstrated that NMPC exerted antimicrobial activity on various phytopathogens of economic relevance in agriculture. According to estimated IC_50_ values, NMPC displayed antifungal action at similar doses on *F. eumartii*, *B. cinerea*, and *P. infestans*. In particular, in *F. eumartii* spores, NMPC drove the cell membrane damage and loss of cell viability. Interestingly, NMPC concentrations needed to reach sublethal doses in *F. eumartii* spores were significantly lower than those previously reported for a chitosan N-derivative (Eweis *et al*, 2006; Liu *et al*, 2018; Wei *et al*, 2019; Zhang *et al*, 2020). In plants, *F. eumartii* infection involves spore attachment and germination before host penetration, lesion formation, and tissue maceration (Prins *et al*, 2000). Thus, our results become of particular importance since the progress of the infection to successfully thrive plant tissues needs the efficient germination of spores and the formation of infective hyphae (Laluk & Mengiste, 2010). *F. eumartii* hyphae proved to be sensitive to NMPC, with an estimated IC_50_ value of 22 μg/mL NMPC. However, *F. eumartii* spore germination registered an IC_50_ much lower (2.5 μg/mL), indicating a cell-specific sensitivity to NMPC. We and others also reported these differential sensitivities of mycelium and spores for other chitosan derivatives (Bautista-Baños *et al*, 2006; Terrile *et al.*, 2015). The spore germination of *F. oxysporum* is more sensitive than hyphal growth to the N-derivative chitosan N-/2(3)-(dodec-2-enyl) succinyl, being the IC_50_ values nearly to 5 μg/mL and 1,000 μg/mL, respectively (Tikhonov *et al*, 2006). A similar effect on *F. oxysporum* spore and mycelial treated with six different quaternary N-alkyl chitosan derivatives was also described (Badawy, 2010). An explanation could be that the lipid membrane composition of fungal cells is a crucial point related to sensitivity to chitosan. Feofilova et al. (2015) reported that linoleic acid predominates in mycelial cells while oleic acid is more abundant in spores of different members of the *Penicillium* genus, supporting the notion of actively growing cellular structures contain more unsaturated lipids than those under exogenous dormancy. In this sense, the ratio between oleic and linoleic acids was higher in conidia than mycelia of *F. oxysporum* and *F. roseum* (Rambo & Bean, 1969). Thus, we hypothesized that the differential sensitivity of NMPC to *F. eumartii* cells (spores and hyphae) is related to the different lipid composition of these cell types. Palma Guerrero et al. (2010) reported that the plasma membrane of other chitosan-sensitive fungi has more polyunsaturated fatty acids than chitosan-resistant fungi. A *Neurospora crassa* mutant with depletion of polyunsaturated fatty acids and reduced membrane fluidity showed increased resistance to chitosan. This finding suggests that cell permeabilization by chitosan may be dependent on membrane fluidity. In both cases, they analyzed the mycelial but not the spore lipid composition.

NMPC has an aminoalkyl phosphonic ligand to which chelating properties (Heras *et al.*, 2001). Ramos et al. (2003) also proved that NMPC is a powerful chelating agent of Ca^2+^ and other ions. Thus, we cannot discard that NMPC as a chelator compound could additionally affect Ca^2+^ levels in *F. eumartii* cells. Kim et al. (2015) demonstrated the Ca^2+^ requirement in fungal developmental processes such as germination, hyphae development, and nutrient uptake (Kim *et al*, 2015). Later, they observed the role of different Ca^2+^ channels in controlling spore germination and hyphal growth in *F. oxysporum* cells (Kim *et al*, 2018). In agreement with this study, the antimicrobial action against *Aspergillus flavus* and *A. parasiticus* correlated with the chelation effect of N-carboxymethyl chitosan by disturbing the uptake of nutritional divalent ions (Cuero *et al*, 1991). However, whether the potent fungicidal action of NMPC involves chelation of Ca^2+^ or other ions needs to be explored.

We showed that NMPC-mediated cell membrane permeabilization was also concomitant with ROS production in fungal spores. Cellular H_2_O_2_ induction may result from the primary effect of cell membrane permeabilization. The ROS production could cause lipids peroxidation of polyunsaturated fatty acids, inducing cell membrane permeabilization (Howlett & Avery, 1997). Both cellular and biochemical events could be responsible for the loss of *F. eumartii* cell viability mediated by NMPC. Similarly to NMPC, induction of cell membrane damage was reported by the chitosan derivatives ATMCS and ATPECS on *F. oxysporum* hyphae (Qin *et al*, 2014). The potential hypothesis that explains the antimicrobial action of chitosan on pathogens relies on the positive charge of the protonated chitosan that enables electrostatic interactions with the negative charging of the pathogen surface hence increasing membrane permeability and subsequently resulting in cell death (Maluin & Hussein, 2020). In this sense, NMPC could also bind to the negative charge of the membrane surface, causing permeabilization in *F. eumartti* spores.

Considering the potential of NMPC as a fungicide, the harmlessness to the plant cells is a crucial feature. Interestingly, lethal doses of NMPC on *F. eumartii* did not exert toxicity in tomato cells. This selectivity was also reported by chitosan and several of their derivatives in different experimental models. Asgari-Targhi et al. (2018) analyzed the effect of bulk or nano-chitosan on the growth and physiology of *Capsicum annuum*. They found that application resulted in non-phytotoxic and extended growth of the seedlings while these effects were dose-dependent. The application of chitosan nanoparticles on wheat and barley showed similar results (Faride *et al*, 2017).

In conclusion, all these findings place NMPC as a very promising eco-friendly biofungicide to protect tomato against different phytopathogens.

## Acknowledgments

This research was supported by the Agencia Nacional de Promoción Científica y Tecnológica (PICT RAICES 0959), Universidad Nacional de Mar del Plata (EXA 928/19), Consejo Nacional de Investigaciones Científicas y Técnicas (CONICET) and Comisión de Investigaciones Científicas (CIC, 1480/18). FAM is a PhD fellow from CONICET. CAC, MCT and MXS are members of the research staff from CONICET. JRM is researcher from CIC. We thank Dr. Candela Lobato and Dr. Ana Laxalt for providing us fungal strains and tomato cells, respectively. We also thank Dr. Rubén D. Conde for his critical reading of our manuscript.

---

**Supplementary figure.**
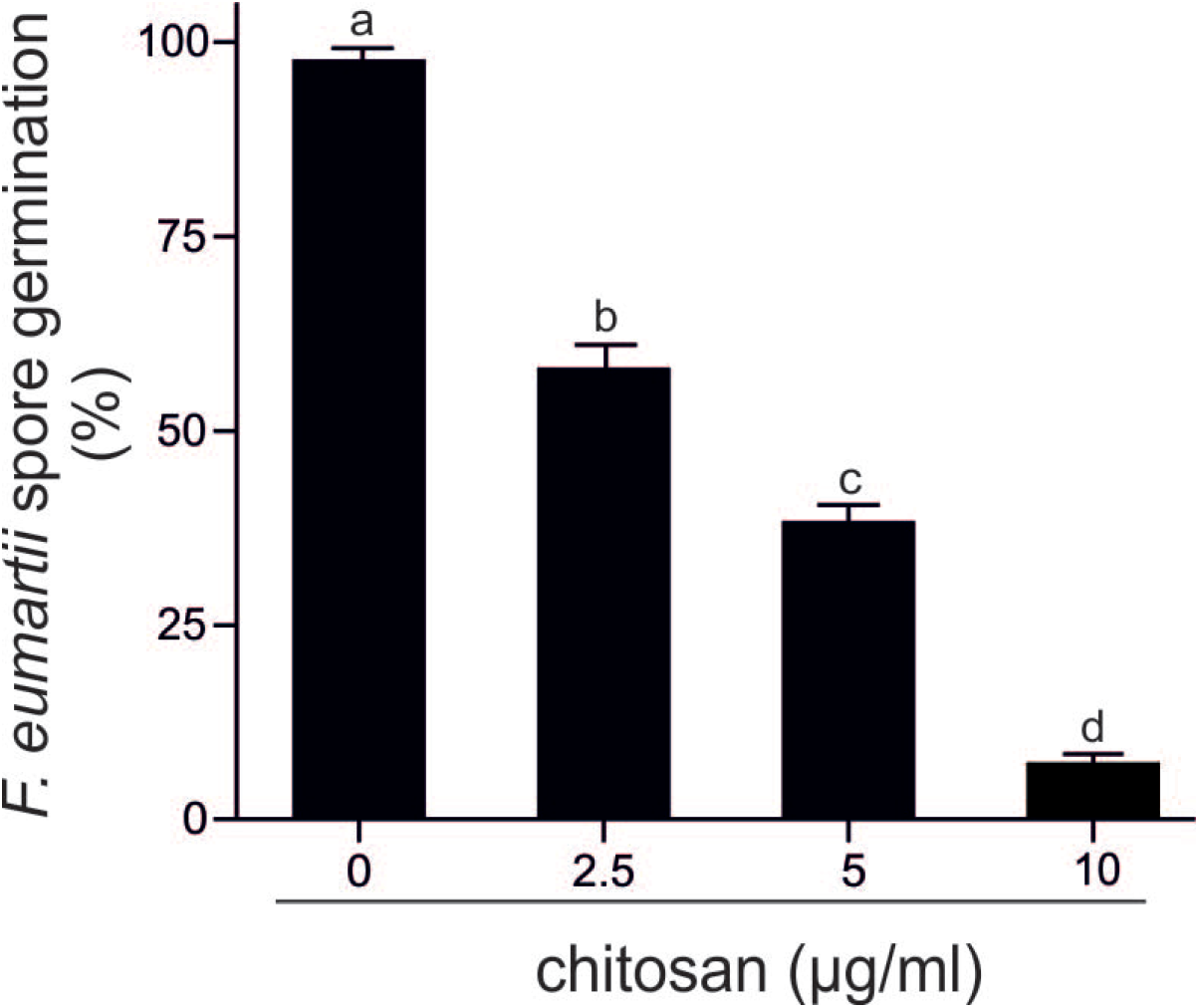
Antimicrobial dose-dependent effect of chitosan. The values represent the percentages of total spores present in each sample after the incubation with different concentrations of chitosan for 24 hr. Each value is the mean ± SD of at least 3 independent experiments.

## References

Andreu AB, Caldiz DO, Forbes GA (2010) Phenotypic expression of resistance to *Phytophthora infestans* in processing potatoes in Argentina. American Journal of Potato Research 87: 177–187

Argüelles-Monal WM, Lizardi-Mendoza J, Fernández-Quiroz D, Recillas-Mota MT, Montiel-Herrera M (2018) Chitosan derivatives: introducing new functionalities with a controlled molecular architecture for innovative materials. Polymers 10: 342

Asgari-Targhi G, Iranbakhsh A, Ardebili ZO (2018) Potential benefits and phytotoxicity of bulk and nano-chitosan on the growth, morphogenesis, physiology, and micropropagation of *Capsicum annuum*. Plant Physiology and Biochemistry 127: 393–402

Badawy M (2010) Structure and antimicrobial activity relationship of quaternary N-alkyl chitosan derivatives against some plant pathogens. Journal of Applied Polymer Science 117: 960–969

Badawy MEI, Rabea EI, Taktak NEM (2014) Antimicrobial and inhibitory enzyme activity of N-(benzyl) and quaternary N-(benzyl) chitosan derivatives on plant pathogens. Carbohydrate Polymers 111: 670–682

Bautista-Baños S, Hernandez-Lauzardo A, Velazquez del Valle MG, Hernandez-Lopez M, Ait Barka E, Bosquez-Molina E, Wilson CL (2006) Chitosan as a potential natural compound to control pre and postharvest diseases of horicultural commodities. Crop Protection 25: 108–118

Benito EP, ten Have A, van ’t Klooster JW, van Kan JAL (1998) Fungal and plant gene expression during synchronized infection of tomato leaves by *Botrytis cinerea*. European Journal of Plant Pathology 104: 207–220

Carpenter CW (1915) Some potato tuber-rots caused by species of Fusarium. Journal of Agricultural Research 5: 183–209

Chirkov SN (2002) The antiviral activity of chitosan. Applied Biochemistry and Microbiology 38: 1–8

Cuero RG, Osuji G, Washington A (1991) N-carboxymethyl chitosan inhibition of aflatoxin production: role of zinc. Biotechnology Letters 13: 441–444

de Oliveira Pedro R, Takaki M, Gorayeb TCC, Bianchi VLD, Thomeo JC, Tiera MJ, de Oliveira Tiera VA (2013) Synthesis, characterization and antifungal activity of quaternary derivatives of chitosan on *Aspergillus flavus*. Microbiological Research 168: 50–55

Deepmala K, Hemantaranjan A, Singh B (2015) Chitosan as a promising natural compound to enhance potential physiological responses in plant: a review. Indian Journal of Plant Physiology 20: 1–9

El Hadrami A, Adam LR, El Hadrami I, Daayf F (2010) Chitosan in plant protection. Mar Drugs 8: 968–987

Eweis M, Elkholy SS, Elsabee MZ (2006) Antifungal efficacy of chitosan and its thiourea derivatives upon the growth of some sugar-beet pathogens. International Journal of Biological Macromolecules 38: 1–8

Faride B, Zeinalabedin TS, Mohamad Zaman K, Seyed Ali Mohamad MS, Ali S (2017) Phytotoxicity of chitosan and SiO_2_ nanoparticles to seed germination of wheat (*Triticum aestivum L.*) and barley (*Hordeum vulgare L.*) plants. Notulae Scientia Biologicae 9

Feofilova EP, Sergeeva YE, Mysyakina IS, Bokareva DA (2015) Lipid composition in cell walls and in mycelial and spore cells of mycelial fungi. Microbiology 84: 170–176

Gupta PK (2018) Toxicity of Fungicides. In: Veterinary Toxicology (Third Edition), Gupta R.C. (ed.) pp. 569–580. Academic Press:

Heras A, Rodríguez NM, Ramos V, Agulló E (2001) N-methylene phosphonic chitosan: a novel soluble derivative. Carbohydrate Polymers 44: 1–8

Hidangmayum A, Dwivedi P, Katiyar D, Hemantaranjan A (2019) Application of chitosan on plant responses with special reference to abiotic stress. Physiology and Molecular Biology of Plants 25: 313–326

Hou Y-P, Qu X-P, Mao X-W, Kuang J, Duan Y-B, Song X-s, Wang J-X, Chen C-J, Zhou M-G (2018) Resistance mechanism of *Fusarium fujikuroi* to phenamacril in the field. Pest Management Science 74: 607–616

Howlett NG, Avery SV (1997) Induction of lipid peroxidation during heavy metal stress in *Saccharomyces cerevisiae* and influence of plasma membrane fatty acid unsaturation. Applied and Environmental Microbiology 63: 2971–2976

Kim H-S, Kim J-E, Frailey D, Nohe A, Duncan R, Czymmek KJ, Kang S (2015) Roles of three *Fusarium oxysporum* calcium ion (Ca^2+^) channels in generating Ca^2+^ signatures and controlling growth. Fungal Genetics and Biology 82: 145–157

Kim H-S, Kim J-E, Son H, Frailey D, Cirino R, Lee Y-W, Duncan R, Czymmek KJ, Kang S (2018) Roles of three *Fusarium graminearum* membrane Ca^2+^ channels in the formation of Ca^2+^ signatures, growth, development, pathogenicity and mycotoxin production. Fungal Genetics and Biology 111: 30–46

Laluk K, Mengiste T (2010) Necrotroph attacks on plants: wanton destruction or covert extortion? The arabidopsis book 8: e0136–e0136

Laxalt AM, Raho N, Have AT, Lamattina L (2007) Nitric oxide is critical for inducing phosphatidic acid accumulation in xylanase-elicited tomato cells. Journal of Biological Chemistry 282: 21160–21168

Liu W, Qin Y, Liu S, Xing R, Yu H, Chen X, Li K, Li P (2018) Synthesis, characterization and antifungal efficacy of chitosan derivatives with triple quaternary ammonium groups. International Journal of Biological Macromolecules 114: 942–949

Malerba M, Cerana R (2018) Recent Advances of Chitosan Applications in Plants. Polymers 10: 118

Maluin FN, Hussein MZ (2020) Chitosan-Based Agronanochemicals as a Sustainable Alternative in Crop Protection. Molecules 25: 1611

Mania S, Partyka K, Pilch J, Augustin E, Cieślik M, Ryl J, Jinn J-R, Wang Y-J, Michałowska A, Tylingo R (2019) Obtaining and characterization of the PLA/chitosan foams with antimicrobial properties achieved by the emulsification combined with the dissolution of chitosan by CO_2_ saturation. Molecules 24: 4532

Mansilla AY, Albertengo L, Rodriguez MS, Debbaudt A, Zuniga A, Casalongue CA (2013) Evidence on antimicrobial properties and mode of action of a chitosan obtained from crustacean exoskeletons on Pseudomonas syringae pv. tomato DC3000. Appl Microbiol Biotechnol 97: 6957–6966

Mendieta JR, Pagano MR, Muñoz FF, Daleo GR, Guevara MG (2006) Antimicrobial activity of potato aspartic proteases (*St*APs) involves membrane permeabilization. Microbiology 152: 2039–2047

Novo DJ, Perlmutter NG, Hunt RH, Shapiro HM (2000) Multiparameter flow cytometric analysis of antibiotic effects on membrane potential, membrane permeability, and bacterial counts of *Staphylococcus aureus* and *Micrococcus luteus*. Antimicrobial Agents and Chemotherapy 44: 827–834

Palma-Guerrero J, Lopez-Jimenez JA, Pérez-Berná AJ, Huang IC, Jansson HB, Salinas J, Villalaín J, Read ND, Lopez-Llorca LV (2010) Membrane fluidity determines sensitivity of filamentous fungi to chitosan. Molecular Microbiology 75: 1021–1032

Plascencia-Jatomea M, Viniegra G, Olayo R, Castillo-Ortega MM, Shirai K (2003) Effect of chitosan and temperature on spore germination of *Aspergillus niger*. Macromolecular Bioscience 3: 582–586

Prins TW, Tudzynski P, von Tiedemann A, Tudzynski B, Ten Have A, Hansen ME, Tenberge K, van Kan JAL (2000) Infection strategies of B*otrytis cinerea* and related necrotrophic pathogens. In: Fungal Pathology, Kronstad J.W. (ed.) pp. 33–64. Springer Netherlands: Dordrecht

Qin Y, Xing R, Liu S, Yu H, Li K, Hu L, Li P (2014) Synthesis and antifungal properties of (4-tolyloxy)-pyrimidyl-α-aminophosphonates chitosan derivatives. International Journal of Biological Macromolecules 63: 83–91

Qiu J, Xu J, Shi J (2014) Molecular characterization of the *Fusarium graminearum* species complex in Eastern China. European Journal of Plant Pathology 139: 811–823

Rambo GW, Bean GA (1969) Fatty acids of the mycelia and conidia of *Fusarium oxysporum* and *Fusarium roseum*. Canadian Journal of Microbiology 15: 967–968

Ramos V, Rodríguez NM, Díaz MF, Rodríguez MS, Heras A, Agulló E (2003) N-methylene phosphonic chitosan. Effect of preparation methods on its properties. Carbohydrate Polymers 52: 39–46.

Rioux D, Jacobi V, Simard M, Hamelin RC (2000) Structural changes of spores of tree fungal pathogens after treatment with the designed antimicrobial peptide D2A21. Canadian Journal of Botany 78 462–471

Romberg MK, Davis RM (2007) Host range and phylogeny of *Fusarium solani* f. sp *eumartii* from potato and tomato in California. Plant Disease 91: 585–592

Sukenik SC, Karuppanan K, Li Q, Lebrilla CB, Nandi S, McDonald KA (2018) Transient recombinant protein production in glycoengineered *Nicotiana benthamiana* cell suspension culture. International Journal of Molecular Sciences 19: 1205

Terrile MC, Mansilla AY, Albertengo L, Rodríguez MS, Casalongué CA (2015) Nitric-oxide-mediated cell death is triggered by chitosan in *Fusarium eumartii* spores. Pest Management Science 71: 668–674

Thordal-Christensen H, Zhang Z, Wei Y, Collinge DB (1997) Subcellular localization of H_2_O_2_ in plants. H_2_O_2_ accumulation in papillae and hypersensitive response during the barley-powdery mildew interaction. The Plant Journal 11: 1187–1194

Tikhonov VE, Stepnova EA, Babak VG, Yamskov IA, Palma-Guerrero J, Jansson H-B, Lopez-Llorca LV, Salinas J, Gerasimenko DV, Avdienko ID (2006) Bactericidal and antifungal activities of a low molecular weight chitosan and its N-/2 (3)-(dodec-2-enyl) succinoyl/-derivatives. Carbohydrate Polymers 64: 66–72

Verlee A, Mincke S, Stevens CV (2017) Recent developments in antibacterial and antifungal chitosan and its derivatives. Carbohydrate Polymers 164: 268–283

Wei L, Tan W, Wang G, Li Q, Dong F, Guo Z (2019) The antioxidant and antifungal activity of chitosan derivatives bearing Schiff bases and quaternary ammonium salts. Carbohydrate Polymers 226: 115256

Younes I, Rinaudo M (2015) Chitin and chitosan preparation from marine sources. Structure, properties and applications. Marine Drugs 13: 1133

Zhang J, Tan W, Li Q, Dong F, Guo Z (2020) Synthesis and characterization of N,N,N-trimethyl-O-(ureidopyridinium)acetyl chitosan derivatives with antioxidant and antifungal activities. Marine Drugs 18: 163

Zhou MG, Wang JX (2001) Study on sensitivity base-line of *Fusarium graminearum* to carbendazim and biological characters of MBC-resistant strains. Acta Phytopathologica Sinica 31: 365–367

Zhu DW, Bo JG, Zhang HL, Liu WG, Leng XG, Song CX, Yin YJ, Song LP, Liu LX, Mei L et al (2007) Synthesis of N-methylene phosphonic chitosan (NMPCS) and its potential as gene carrier. Chinese Chemical Letters 18: 1407–1410

